# Identifying priority conservation areas for recovering large carnivores using citizen science data

**DOI:** 10.1101/454603

**Authors:** A-S. Bonnet-Lebrun, A.A. Karamanlidis, M. de Gabriel Hernando, I. Renner, O. Gimenez

## Abstract

Understanding the processes related to wildlife recoveries is not only essential in solving human – wildlife conflicts, but also for identifying priority conservation areas and in turn, for effective conservation planning. We used data from a large citizen science program to study the spatial processes related to the demographic and genetic recovery of brown bears in Greece and to identify new areas for their conservation. This was achieved by visually comparing our data with an estimation of the past distribution of brown bears in Greece and by using a Point Process Model to model habitat suitability, and then comparing our results with the current distribution of brown bear records and with that of protected areas. Our results indicate that in the last 15 years bears may have increased their range by as much as 100%, by occupying mainly anthropogenic landscapes and areas with suitable habitat that are currently not legally protected, thus creating a new conservation reality for the species in Greece. This development dictates the re-evaluation of the national management and conservation priorities for brown bears in Greece by focusing in establishing new protected areas that will safeguard their recovery. Our conservation approach is a swift and cheap way of identifying priority conservation areas, while gaining important insights on the spatial processes associated with population recoveries. It will help prioritize conservation actions for brown bears in Greece and may serve as a model conservation approach to countries facing financial and logistic constraints in the monitoring of local biodiversity or facing challenges in managing rapid population recoveries. Our conservation approach appeared also to be better suited to identifying priority areas for conservation in areas with recovering wildlife populations and may therefore be used as an “early-warning” conservation system.

## Introduction

Large carnivores have been celebrating significant population comebacks in Europe (Chapron *et al.*, 2014), sparking an increased interest in identifying the preconditions, factors and processes that have facilitated these comebacks (e.g. López-Bao *et al.*, 2015). A thorough understanding of the processes related to large carnivore recoveries is not only essential in solving the increasing number of human – large carnivore conflicts (Bautista *et al.*, 2016), but also for predicting potential habitat in areas of large carnivore population recovery and in turn, for effective conservation planning in the European context of human-dominated landscapes (Linnell *et al.*, 2008). At the same time, the vulnerability to human threats (Ripple *et al.*, 2014), the socio-cultural and financial implications of human – wildlife conflicts (Treves & Karanth, 2003) and the often large spatial requirements (Ripple *et al.*, 2014, Newsome & Ripple, 2015) and long dispersal abilities (Kojola *et al.*, 2006) of large carnivores dictate a management and conservation approach that is based on the swift collection and evaluation of accurate data on their status. This ensures their recovery and survival through their adequate representation inside and outside protected areas (Di Minin *et al.*, 2016).

As new technologies emerge that make it easier to collect and transmit information on one’s location, volunteer participation in data collection is becoming increasingly more important as an ecological research tool (Dickinson *et al.*, 2012). Volunteers participating in citizen science programs can collect more data and cover wider areas, faster than researchers alone would, all of this at a lower cost (Dickinson *et al.*, 2010, Dickinson *et al.*, 2012). If analysed properly, opportunistic, presence-only data of citizen science projects may produce reliable estimates of wildlife distribution trends (van Strien *et al.*, 2013); however, such data may suffer from *observer bias* [*sensu* Warton et al. (2013), also commonly referred to as *sampling bias*], when the sampling effort is not evenly distributed in the entire study area. Thus, if a species is present, it is more likely to be recorded when more people are present to see it, i.e. in areas more densely populated and/or more accessible, such as those close to roads, research centres and cities (Geldmann *et al.*, 2016, Daru *et al.*, 2018). If no correction is made for observer bias, the risk is to model the observer distribution instead of the species distribution (Hortal *et al.*, 2008, Warton *et al.*, 2013). To address this issue, Warton et al. (2013) developed a method that corrects for observer bias by relating the records of species presence to variables that are split into two categories: *ecological variables* are variables that are likely to influence a species’ occurrence, whereas *observer bias variables* are variables that are likely to influence the detection of a species (the observer’s occurrence mainly). When making predictions, only the ecological variables are used, and a common value for the observer bias variables (e.g. the average, or the minimum/maximum value), making the predictions as if the distribution of observers was spatially homogeneous. This method has been successfully applied to model the distribution of plant data (Warton *et al.*, 2013), but is yet to be applied to animals.

Presence-only data obtained from citizen science programs are particularly relevant to large carnivores, such as brown bears (*Ursus arctos*), because bears are difficult to monitor, due to their cryptic and solitary nature and due to their relatively low density of occurrence over large areas (Kindberg *et al.*, 2009). At the same time however, bears are also highly charismatic species attracting public attention, increasing the chances that observations would be reported, thus making them particularly suitable for citizen science programs.

Brown bears are globally considered by the IUCN as species of Least Concern. In Europe several populations are small, isolated and threatened by habitat loss and fragmentation and by human - bear conflicts (Swenson & Sandegren, 2000, Bautista *et al.*, 2016, Piédallu *et al.*, 2017). This is particularly the case for brown bears in Greece where the species reaches its southernmost European distribution and is considered to be endangered, numbering fewer than 500 individuals (Karamanlidis *et al.*, 2015). Despite increasing human - wildlife conflicts (Karamanlidis *et al.*, 2011), bears have been recovering in recent years (i.e. after approximately the year 2000) in Greece, both demographically (Karamanlidis *et al.*, 2015) and genetically (Karamanlidis *et al.*, 2018). At the same time, circumstantial evidence suggests that the species has also been expanding its range (Karamanlidis *et al.*, 2008); however, no thorough, nation-wide study has been conducted so far to substantiate this fact, partially because of the logistic and financial constraints that have befallen the country since the onset in 2009 of a financial crisis. This incomplete understanding of the current distribution of brown bears in Greece is hindering their effective conservation, which in turn may compromise the ongoing demographic and genetic recovery of the species in the country.

The aims of this study were to take advantage of a large citizen science data set to study the presence of brown bears in Greece during the recovery of their population in the country and to identify new priority areas for their conservation. The first aim was achieved by visually comparing our citizen science data with an estimation of the past distribution of the species in the country. The latter aim was achieved by applying Warton et al.’s (2013) approach to our citizen science data to model habitat suitability, and then comparing the predictions of our model with the current distribution of brown bear records in Greece and with that of protected areas in the country. This comparison allowed us to assess the distribution of highly suitable bear habitat in relation to the distribution of bears and protected areas and obtain new, valuable insights on the spatial recovery and conservation priorities of brown bears in Greece.

## Materials and methods

### Data collection

Data on bear presence were collected from 2004 – 2016 within the framework of a citizen science program, the “Hellenic Brown Bear Rescue and Information Network” (HBBRIN), established by the non-governmental organization ARCTUROS. Information on bear presence (i.e. opportunistic observations, damage to human property, attacks to humans, approaches to inhabited areas) was received through the post, telephone or email and was verified either on site by a field team or through the evaluation of the information provided (e.g. photographs and videos).

### Studying brown bear presence in Greece

In order to gain insights on the spatial processes during the demographic and genetic recovery of brown bears in Greece, we mapped bear records from the HBBRIN using QGIS v2.14 (QGIS Development Team, 2016) and compared them with the past distribution of the species in the country. We considered as a reference the only account of the distribution of brown bears in Greece (Fig. 1) published in the assessment of the species for the Red Book of Endangered Species of Greece (Mertzanis *et al.*, 2009). In this assessment the total area of continuous brown bear range in Greece was estimated at approximately 13,500 km^2^, which consisted of two geographically distinct population nuclei in the northeastern and northwestern part of the country. Although no information is available on how this map has been produced and whether it represents only core areas of the species or also areas of temporal re-occurrence, it is evident, from the references therein and its publication date, that the map is the best available account on the distribution of brown bears in Greece prior to their demographic and genetic recovery and the beginning of our study.

**Figure 1.**
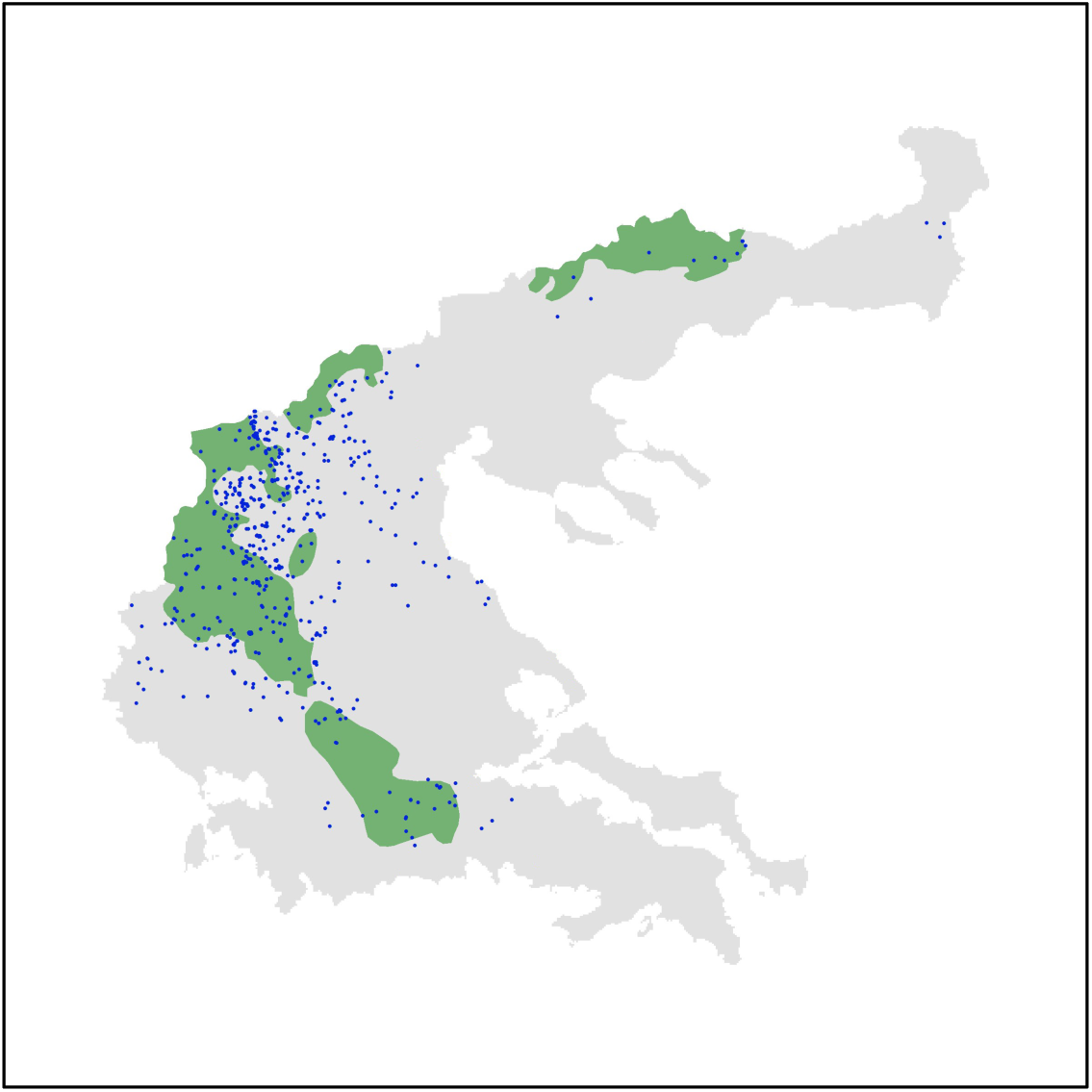
Map of a part of continental Greece indicating the distribution of brown bears in the country according to Mertzanis *et al.* 2009 (shaded green area) and the locations of 632 brown bear occurrences recorded through the HBBRIN (2004 – 2016).

In order to compare the habitat characteristics between the past distribution of brown bears in Greece and the new areas where the species was documented through the HBBRIN, we calculated the percentage of coverage of different habitat types [modified from Corine Land Cover 2006 (CLC) seamless vector data, Version 16 (European Commission, 04/2012)], road density (OpenStreetMap, 2016) and population density (Center for International Earth Science Information Network (CIESIN) & Centro Internacional de Agricultura Tropical (CIAT), 2005). The size of the area where the species was documented through the HBBRIN was calculated by Local Convex Hulls (LoCoHs) using a fixed radius of 50 km and the five nearest neighbors of each point (R package “rhr”; (Signer, 2016), excluding however 5% of the outermost records (i.e. with respect to the past distribution area) in order to reduce the impact of extreme observations.

### Identifying new priority areas for brown bear conservation in Greece

#### Modelling framework

We modelled the intensity of observations per space unit as our response variable in a Point Process Model (PPM), a common tool for modelling the occurrence of species based on a set of explanatory factors using presence-only data (Renner *et al.*, 2015; see Appendix 1). Using the method by Warton et al. (2013), the fitted PPM produces estimates of the intensity of brown bear observations per square kilometre, which allows us to infer habitat suitability based on a set of ecological variables after correcting for observer bias variables. By correcting for observer bias, the output is detrended and hence can be interpreted as an intensity of bear observations per square kilometre, if all locations in the study area had equal sampling effort as quantified by the observer bias variables (Warton *et al.*, 2013). If we can accept that the realised distribution of brown bears in Greece is compatible with preferred habitat, then such maps of corrected intensity are proportional to habitat suitability. After initially fitting an inhomogeneous PPM to the brown bear records, we found that there was additional clustering of the records beyond that which was explained by the model. Therefore, we used an alternative PPM which accounts for spatial dependence among point locations, an area-interaction model (see Appendix 1).

#### Explanatory variable selection

A grid of 1×1 km^2^ was laid over the study area, which corresponds to the resolution of the coarsest explanatory variable. The mean value of a set of explanatory variables expected to influence either bear or observer presence was calculated for each grid cell. Explanatory variables were chosen following previous studies of brown bear in Europe (Naves *et al.*, 2003, Martin *et al.*, 2013). We used topography (i.e. maximum altitude and mean slope), distance to forest edge and distance to shrubland edge, density of rivers (i.e. accumulated length of rivers in the pixel), and percentage of agricultural land as ecological variables, and distance to human settlements, as well as distance to the closest primary or secondary road as observer bias variables (Appendix 1, Table S1). The construction of the grid and the calculation and extraction of the ecological variables were done in the R software (R Development Core Team, 2011) and in QGIS (Quantum GIS Development Team, 2013).

#### Comparison of habitat suitability predictions with bear records and protected area distribution in Greece

The Natura 2000 (N2K) network is a European network of protected areas for birds and habitats (http://ec.europa.eu/environment/nature/natura2000/db_gis/index_en.htm). Some of the sites in the N2K network (33 out of 419) are included in the dedicated network for the protection of bears in Greece [as reported in their Standard Data Forms (SDFs)]. To compare our habitat suitability predictions with the presence of bears and the distribution of protected areas in Greece, we mapped all the N2K sites in Greece (http://www.ekby.gr/ekby/en/Natura2000_main_en.html; last updated in 2012), identified the sites included in the dedicated network for the protection of bears in Greece and those which were not. For this, we mapped only areas of highly suitable habitat, defined as those grid cells with predicted intensities above the 80% quantile. Only N2K sites with more than 5% of their total area classified as highly suitable habitat were retained. Areas containing highly suitable habitat, but not included in the N2K network of protected areas, were also identified.

## Results

### Studying brown bear presence in Greece

From 2004 – 2016 the HBBRIN recorded the presence of bears on 632 occasions: 40% of these cases (i.e. 251 data points) were located within the past distribution of the species, 60% beyond (Fig. 1). The new distribution area of brown bears in Greece that was not included in the map of Mertzanis et al. (2009) covered a total of 16,661 km^2^ and was, compared to the past distribution of the species, characterized by lower elevations, higher coverage by primary and secondary roads and agricultural areas and lower coverage by mature forests (Table 1).

**Table 1.**
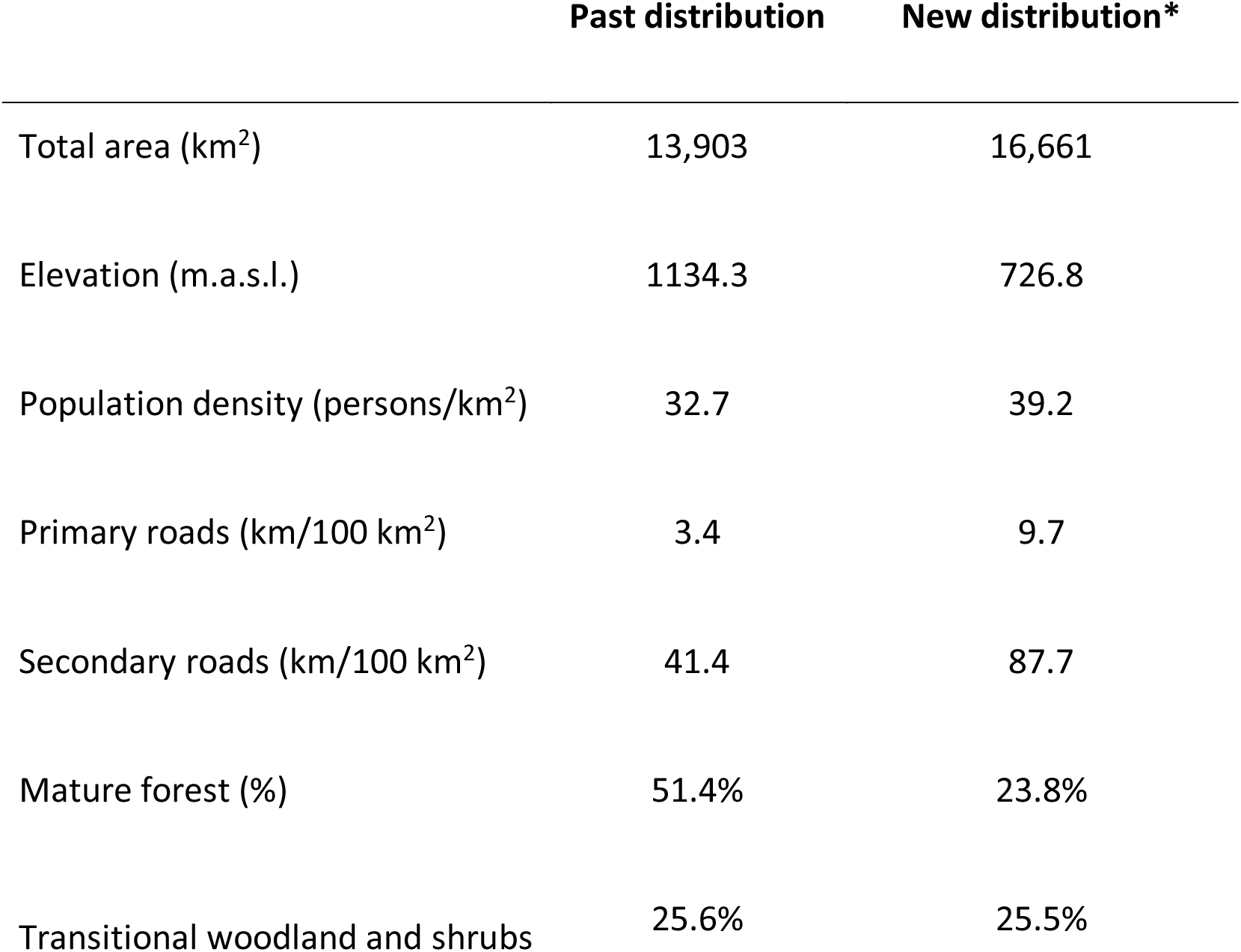
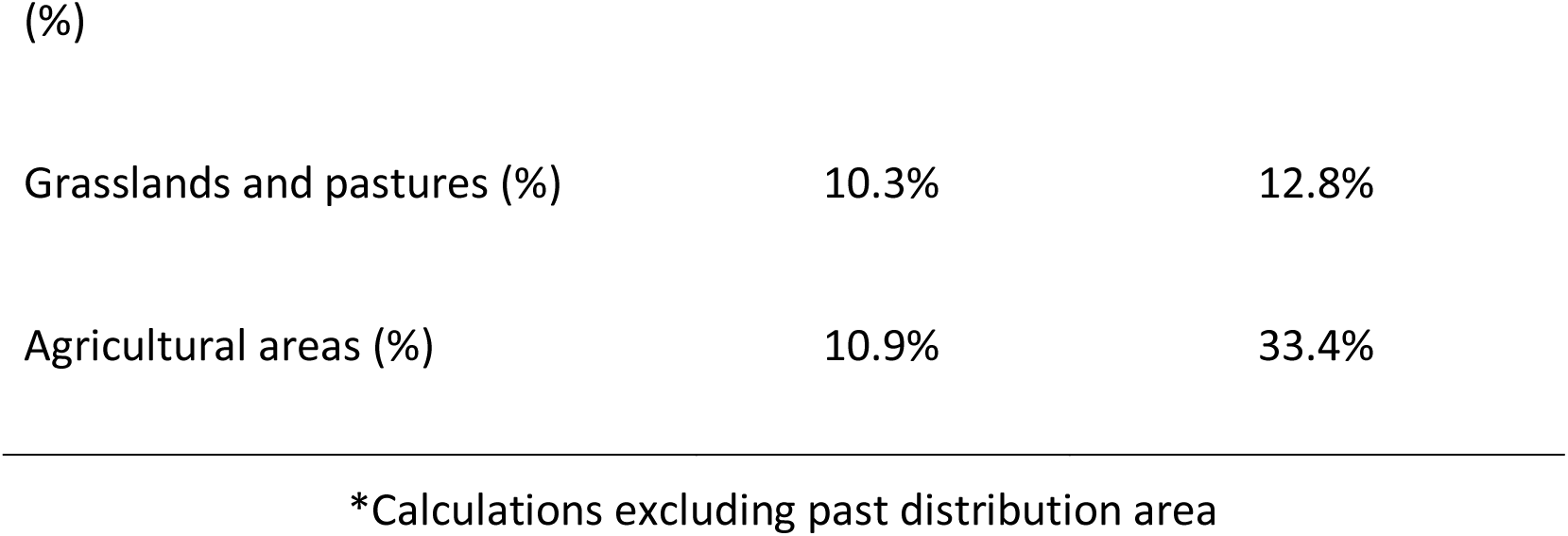
Habitat characteristics of the past and new distribution of brown bears in Greece (for more information, see the “Studying brown bear presence in Greece” in the Materials and methods section).

|  | Past distribution | New distribution* |
| --- | --- | --- |
| Total area (km<sup>2</sup>) | 13,903 | 16,661 |
| Elevation (m.a.s.l.) | 1134.3 | 726.8 |
| Population density (persons/km<sup>2</sup>) | 32.7 | 39.2 |
| Primary roads (km/100 km<sup>2</sup>) | 3.4 | 9.7 |
| Secondary roads (km/100 km<sup>2</sup>) | 41.4 | 87.7 |
| Mature forest (%) | 51.4% | 23.8% |
| Transitional woodland and shrubs | 25.6% | 25.5% |
| Grasslands and pastures (%) | 10.3% | 12.8% |
| Agricultural areas (%) | 10.9% | 33.4% |

\*Calculations excluding past distribution area

### Identifying new priority areas for brown bear conservation in Greece

We first assessed the effect of each covariate on the intensity of brown bears per square kilometre as estimated in the PPM. To visualise these effects, we produced scatterplots of the predicted intensities vs. standardised covariates, and then fitted a smoothed generalized additive model to produce a response curve (Fig. 2). The intensity of the PPM was the highest for intermediate altitudes, slopes and percentage of agricultural land (Fig. 2, Appendix 1, Table S2). The intensity of the PPM decreased for greater distance to shrubland and forest edges and to both types of roads and human settlements (Fig. 2, Appendix 1, Table S2).

**Figure 2.**
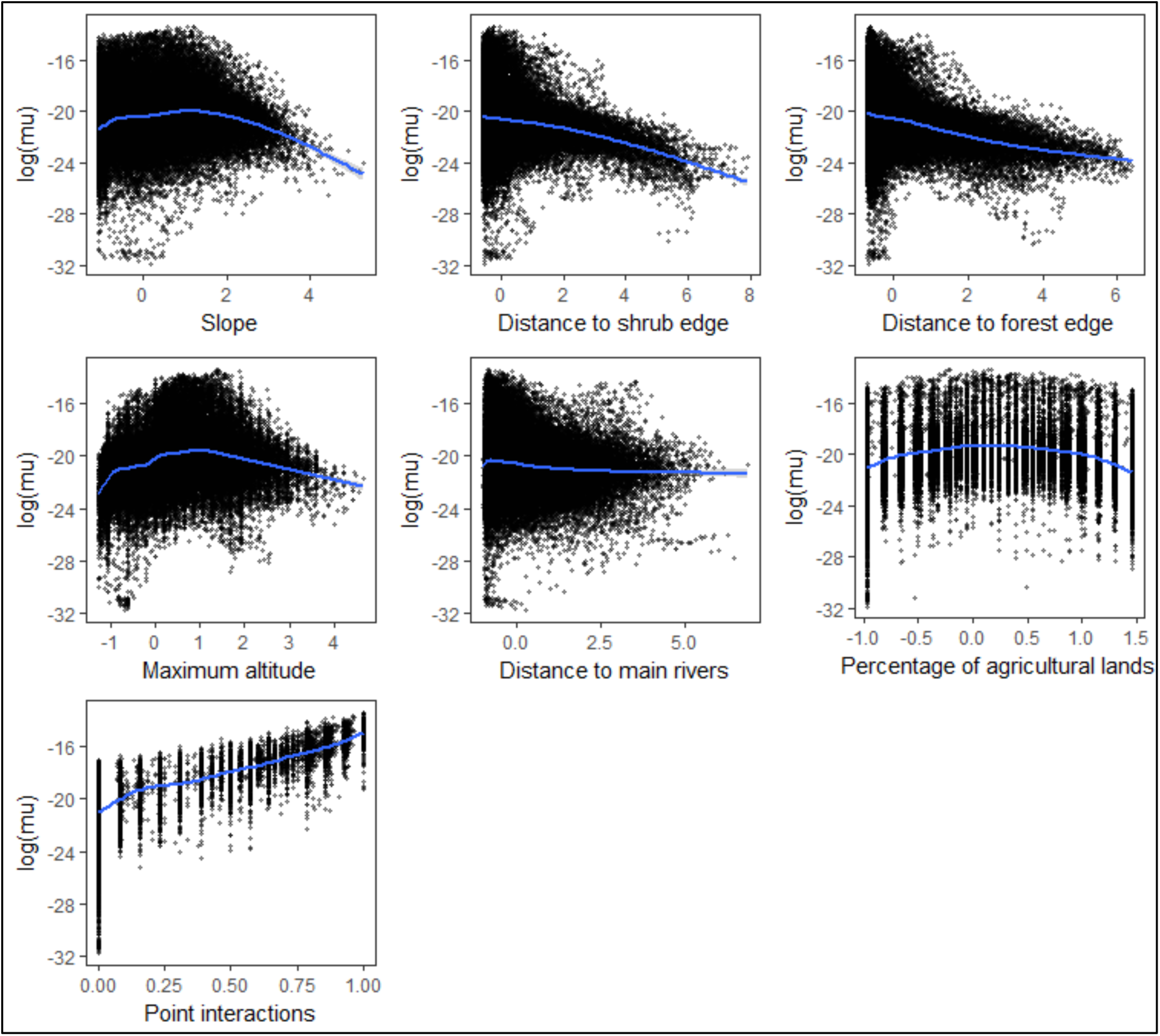
Response curves for each environmental variable in the model. Points: scatterplot of predicted intensity against standardised covariate values. Blue line: fitted smooths (using generalised additive models).

When examining the predictions from the model (Fig. 3), habitat suitability was patchier when conditioning on a common level of bias (Fig. 3A: best model, correcting for observer bias) than when *observer bias* was ignored (Fig. 3B: only *ecological* variables were used).

**Figure 3.**
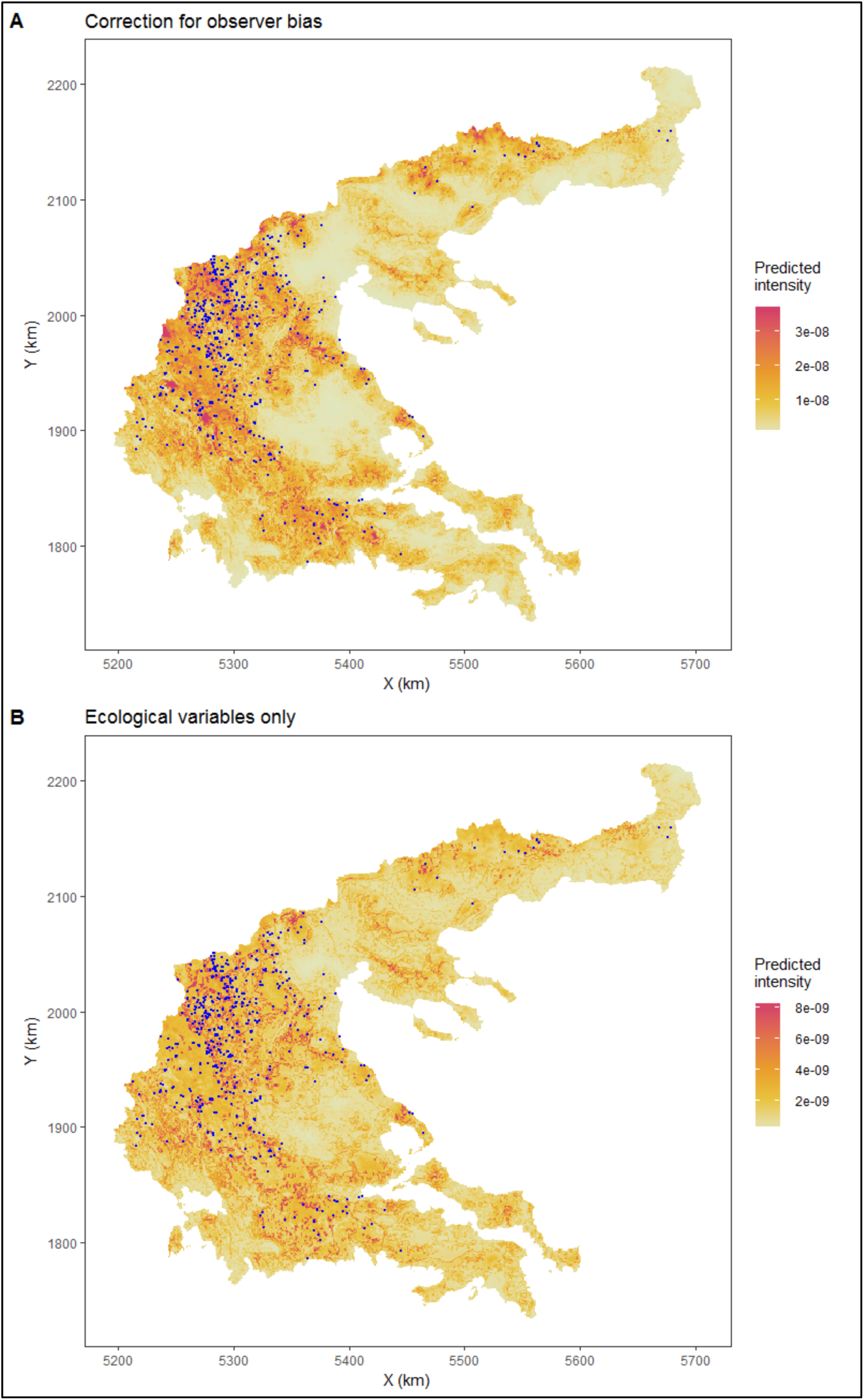
Habitat suitability map from the model with point interaction, A: from the model with both ecological and observer bias variables, where predictions were conditioned on a common level of bias; B: from the model that did not correct for observer bias (i.e. with the ecological variables only); blue points are the opportunistic observations. For visualisation purposes, values above the 0.95 percentile of the global map are set to the 0.95 percentile value.

Our habitat suitability analysis identified 99 N2K sites with highly suitable brown bear habitat in Greece (i.e. predicted intensities higher than the 80% quantile covering more than 5% of the total area of an N2K site) (Fig. 4). Twelve N2K sites contained > 80% of highly suitable habitat, 25 contained > 60%, 53 contained > 40%, while the remaining N2K sites contained < 40% of suitable brown bear habitat respectively (Appendix S2). 33 of the N2K sites with suitable bear habitat in Greece had bears included in their SDFs, 66 did not. N2K sites with bears in their SDFs were associated with high-quality bear habitat (88% of these sites had > 40% suitable bear habitat) and actual bear presence (in 64% of these sites, 81 cases of bear presence were recorded). N2K areas without bears in their SDFs were associated with lower-quality bear habitat (36% of these sites had > 40% suitable bear habitat) and lower bear presence (in 45% of these sites, 104 cases of bear presence were recorded). There were however notable exceptions, including 7 areas with high quality bear habitat (i.e. > 60%) and bear presence, as well as 5 areas with low quality bear habitat (i.e. < 20%) and bear presence (Appendix S2). In addition, bear presence was recorded 485 times in areas not included at all in the N2K network (Fig. 4), mainly in the western population nucleus of the species in the country. High-quality habitat in this part of the range of the species was located mainly in the area enclosed by the Grammos, Vitsi-Varnoundas and Askio mountains (Fig. 4).

**Figure 4.**
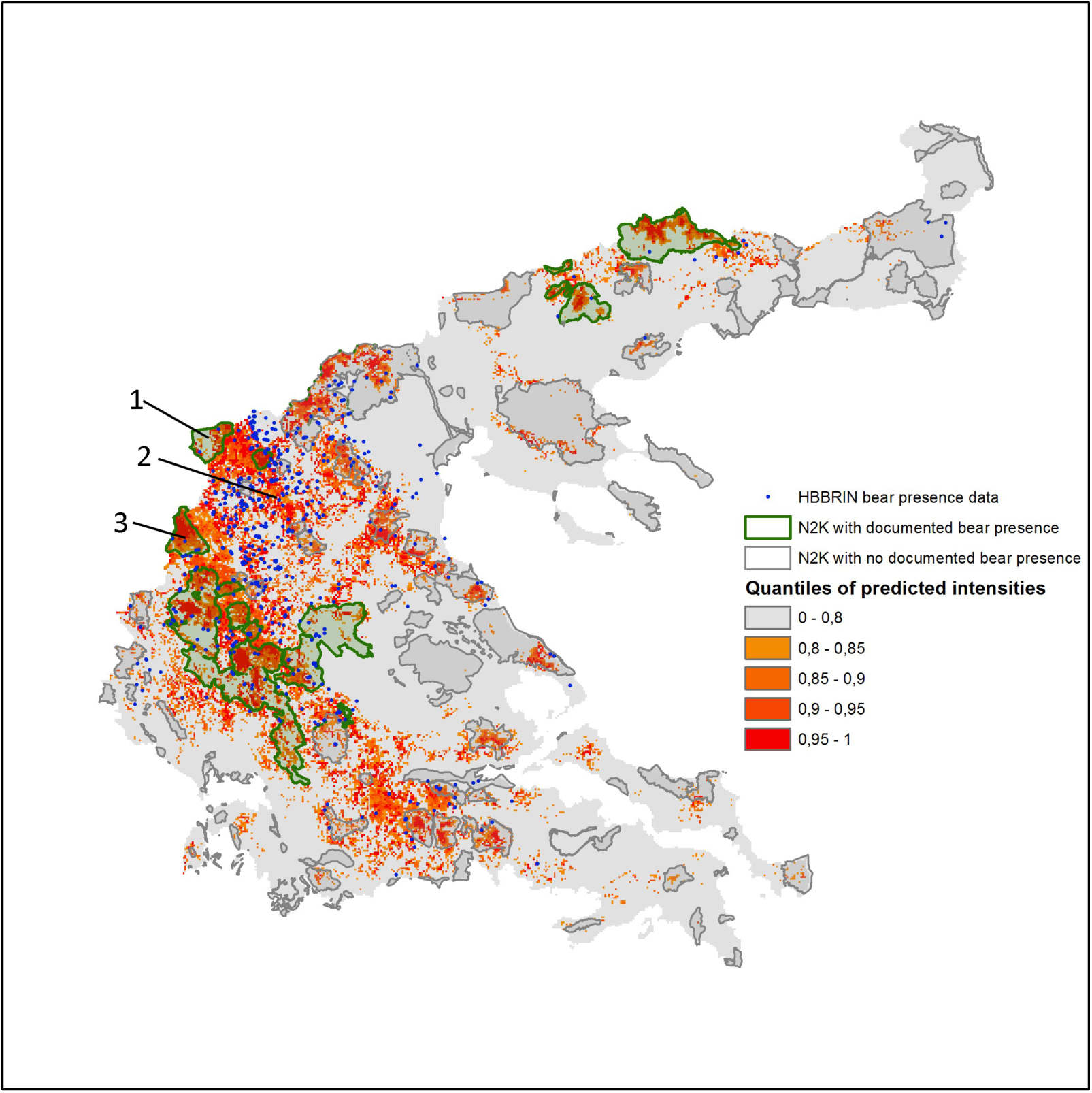
Map of a part of continental Greece indicating the opportunistic data (blue), the predicted highly suitable habitat (red gradient) and the Natura 2000 (N2K) coverage with (green) and without presence of bears (gray) in SDFs. Habitat suitability predictions were obtained with the model with point interactions, built with both ecological and observer bias variables, and projected conditioning on a common level of bias. All grid cells for which predicted intensity was above the 0.80 percentile of the global map qualified as highly suitable habitat. Numbers indicate the Mountains: 1) Vitsi-Varnoundas, 2) Askio, and 3) Grammos, respectively.

## Discussion

Understanding the processes associated with population recoveries and identifying areas that are suitable for recolonization by endangered species is essential to support effective conservation policies (Cianfrani *et al.*, 2010). We used data from a citizen science project and habitat suitability modelling to understand the spatial processes associated with the demographic and genetic recovery of an endangered brown bear population at the southern edge of its European distribution and to identify priority areas for conservation.

Brown bear presence during the demographic and genetic recovery of the species in Greece was recorded in both the northeastern and northwestern nuclei of the species in the country. However, a significant percentage of the bear records of the study fell outside of the past distribution of the species in Greece, confirming previous circumstantial evidence (Karamanlidis *et al.*, 2008) and providing for the first time a clear indication of the spatial recovery of the species and the recolonization of new areas. This spatial recovery has been associated primarily with the recolonization of anthropogenic/cultural landscapes (Table 1) and is consistent with the increase in human – bear conflicts (Karamanlidis *et al.*, 2011) and in bear – vehicle collisions (Karamanlidis *et al.*, 2012) recorded in Greece during this time. It is also consistent with our understanding of the genetic recovery of the species in the country (Karamanlidis *et al.*, 2018) and our understanding of the effects of human activity on the activity and habitat selection patterns of brown bears in Greece (de Gabriel Hernando et al. In Review). Although distribution data from the HBBRIN should be viewed with caution, as this is a citizen science project based in the western part of Greece (and therefore more popular in this part of the country) mainly related to the provision of expertise assistance in human – bear conflicts and cases of bears in distress, evidence suggests that bears in Greece in the last 15 years may have increased their range by as much as 100%. Further research is necessary to substantiate this fact and complete our understanding of the spatial recovery of this large carnivore in Greece.

The results of the study indicate that bear habitat suitability decreased further away from the forest and from the shrubland edge (Fig. 2). We interpret the first effect as forest and shrubland edges providing an interface between potential resources outside and refuges inside. Forest cover in itself (which can be interpreted as providing refuge) has already been shown to be an important variable influencing bear presence (Naves *et al.*, 2003, Martin *et al.*, 2012). The quadratic effects found for altitude (consistent with Güthlin *et al.*, 2011), slope and percentage of agricultural land could be also due to the interface between refuge (i.e. higher altitude, steeper slopes and fewer open spaces) and potential resources (i.e. lower altitudes, smooth terrain and existence of crop fields).

Although previous studies have tried to correct for some of the spatial sampling bias inherent to citizen science data (e.g. Phillips *et al.*, 2009), these approaches would not have been suitable for our case study. For example, volunteers often record within a given study area several other species, usually more common than the focus species, providing information that can be used to infer observer bias and correct for it (e.g. Phillips *et al.*, 2009). However, this approach merely replaces observer bias with species richness bias (Warton *et al.*, 2013), because there might be spatial patterns in the distribution of species richness. Also, this approach would not be applicable to bears and large carnivores in general, because it is hard to find species that would be likely to be recorded by the same citizens at the time of the observations. Finally, in cases where, like in our study, citizens spontaneously report opportunistic data, it is almost impossible to rely on any type of information other than the location of the observations. By relying on independently collected data to model observer bias, the method proposed by Warton et al. (2013) has the advantage to provide a flexible and generic way to correct for observer bias.

However, several issues arise when trying to model observer bias using this approach. First, one needs to select variables that capture observer bias. In doing so, some variables seem intuitive, like those related to human population density or to accessibility of the area. But others can be harder to classify, like, for example, distance to human settlements, which may have a negative effect on species detection with the less people in the area, the less likely that someone will detect the species if it is present. Therefore, it seems plausible to consider distance from human settlements as an *observer bias* variable. This is likely to be the case for plants, but for free-ranging mammals with large home ranges, the same variable can also have two opposite ecological effects, with bears either avoiding populated areas (e.g. Naves *et al.*, 2003, Martin *et al.*, 2012, Piédallu *et al.*, 2017) because they are perceived as risky, or being attracted to them, as they can provide shelter and/or resources (Elfström *et al.*, 2014). Such difficulty of classifying variables can create limitations because if some features appear both in the ecological and in the observation process, the model is non-identifiable (Fithian & Hastie, 2013). Similar problems of joint effects on presence and detectability can arise with roads, which can provide access for observers to the sites where bears are present (*observer bias* variable), but can also be related to disturbance (habitat fragmentation, road mortality) and thus negatively influence the presence of bears (Graves *et al.*, 2011, Martin *et al.*, 2012). We chose to consider the distance to roads as an *observer bias* variable. Although its negative effect on bear detection (i.e. the further away from a road, the less likely a bear observation will be made) is consistent with that choice, the difficulty to decide *a priori* whether a variable should be considered as *observer* or *ecological bias* emphasizes the need for future research.

Ideally, one should validate the approach – at least for a subset of the study area – using data collected by standardised protocols (hence not suffering from observer bias). This would allow concluding whether the correction for observer bias improved the predictions or not, and would also allow for a model which includes both presence-only and presence-absence data. However, in this study we did not have access to such good-quality validation data. We had access to telemetry data (VHF and GPS data – which are commonly available for a subset of the study area in many projects), but using these for validation would not have been appropriate for several reasons. First, telemetry data are biased towards the capture locations – it is impossible to assign areas with no records to areas without the species or to areas with the species but where no tracked individual occurred. Similarly, areas with a lot of telemetry locations could correspond to areas that are preferred by the bears or simply to areas where the home range of multiple tracked individuals overlap. A solution to this problem would be to model habitat selection within each individual’s home range. However, the habitat selection that would be modelled using this approach is at a scale that does not match the habitat selection modelled using Warton et al.’s method (2013) for opportunistic data. The former is Johnson’s *first order selection*, while the latter is *third order selection* (Johnson, 1980). In particular, modelling selection within the range does not inform on which environmental conditions are highly unsuitable for the species (by definition, no telemetry data will come from these areas), whereas this distinction between suitable and unsuitable conditions is what is the most important for conservation. This distinction is also where the strength of our predictions lie (i.e. identifying areas that are highly suitable vs. areas that are not – whereas areas with mixed predictions are less useful). Overall, despite the fact that it was not possible to formally validate our correction for observer bias, studies that have had access to good validation datasets have already demonstrated the added value of correcting for observer bias using the approach we used here (e.g. Warton *et al.*, 2013). This is supported in our study also by the findings presented in Fig. 4 (i.e. comparison between our predictions and N2K sites with known bear presence), which indicate that most sites with known presence of bears are predicted as containing highly suitable habitat.

Apart from being generally applicable to species that are rare and sampled by citizen science and for which there is only presence-only data, our use of Warton et al.’s method (2013), is of important, practical relevance for (endangered) species conservation in general and brown bear conservation in Greece in particular.

Our habitat suitability modelling has produced a map with the predictions of the most suitable bear habitat in the country. The predictions seem less accurate in the northeastern nucleus where a high percentage of observations fall outside areas with high predicted intensities. This could be due to the existence of ecological differences between the Pindus and Rodopi bear populations and the fact that only a 2% of the total observations used for modelling come from the latter, thus predicting habitat suitability with less accuracy in that nucleus. As evidenced by these predictions, the existing network of protected areas that is formally dedicated to the protection of the species (i.e. N2K sites with bears included in their SDFs) contains critical habitat for the conservation of the brown bear in Greece. This fact is indirectly supported by the high densities of bears (Karamanlidis *et al.*, 2015), the high number of human – bear conflicts (Karamanlidis *et al.*, 2011) and the genetic importance as source populations in these areas (Karamanlidis *et al.*, 2018) and therefore these areas should be considered as belonging currently to the core habitat of brown bears in Greece. However, the number of bear observations collected through the HBBRIN in these areas does not correspond to their formal importance as expressed through their legal protection, as only 12.3% of bear observations from the HBBRIN were recorded there. This discrepancy is likely explained by the increased experience (Galloway *et al.*, 2006) of people living in areas with well-established bear populations, as opposed to people living in areas where bear populations are recovering and bear presence is still an unusual event. This suggests furthermore that our approach might be better suited to identifying priority areas for conservation in areas with recovering wildlife populations, i.e. may be used as an “early-warning” conservation system.

The results of our study indicate furthermore that only a portion of highly suitable habitat for bears in Greece is currently included in the dedicated N2K network of protected areas for bears; most records of bears originated either from areas with no legal protection status or from protected areas with no specific management measures for the protection of bears, thus creating a new “conservation reality” for the species in the country. On a practical level, this new conservation reality dictates that protected areas that do not have the brown bear in their SDFs, but do have highly suitable habitat and bear presence at the same time in them, will need to prepare for the possibility of the re-establishment of the species in their management area and adjust their management priorities and actions accordingly. For areas with highly suitable habitat and bear presence that are not legally protected, this conservation reality dictates that the information of this study should be used by the national conservation authorities to re-evaluate their national management and conservation priorities, while focusing at the same time in establishing new protected areas for the species. A similar approach has been suggested for the conservation of another recovering large carnivore in Greece, the grey wolf (*Canis lupus*) (Votsi *et al.*, 2016).

The main differences in habitat characteristics between the past and new distribution of brown bears in Greece provide insights into the spatial recovery of the species; new distribution areas are more humanized (i.e. denser road network, higher population densities, higher proportion of agricultural areas and lower proportion of mature forest). The ability for bears to re-colonize such areas is most likely explained by multiple factors including: 1) the behavioural plasticity of the brown bear (Ordiz *et al.*, 2014), which in this case has resulted in activity and habitat selection adaptations to human activity of bears in Greece (de Gabriel Hernando et al. In Review); and (2) abandonment or decrease in agricultural activities in the less productive areas as a consequence of the general rural abandonment, allowing a progressive naturalization of these areas (Poyatos *et al.*, 2003).

## Conclusions

For the past several years Greece has suffered a financial crisis that has had a negative effect on the national environmental management apparatus (Lekakis & Kousis, 2013) that is likely to leave national management authorities in the future struggling to find the necessary funds to effectively monitor and manage biodiversity in the country. Given that the bear population in the country is rapidly recovering (Karamanlidis *et al.*, 2015, Karamanlidis *et al.*, 2018) and that at the same time bear densities and conflicts with humans in Greece are increasing (Karamanlidis *et al.*, 2011), our approach of collecting information on bear presence through a citizen science program and using it to produce habitat suitability maps has been a swift and cheap way of identifying potential hot-spots of bear presence, activity and conflicts with humans in the country, while gaining important insights on the spatial processes associated with the recovery of this large carnivore.

Our approach will help prioritize conservation actions in the country towards the areas that need it the most and serve as a model approach to other countries facing similar financial and logistic constraints in the monitoring of local biodiversity or facing similar challenges in managing the rapid recovery of a large carnivore. Acknowledging the previous, we propose the intensification of efforts in Greece to further develop the Hellenic Brown Bear Rescue and Information Network, by carrying out targeted awareness campaigns to the general public and selected stakeholders (e.g. Forestry and Veterinary Departments, Management Authorities of Protected areas) that will increase data input and ultimately the quality of the habitat suitability maps produced using this method. At the same time, it is clear that in the case of the brown bear the current setup of the protected areas network in Greece does not reflect the current conservation reality and that there is a clear need to re-evaluate the existing network of protected areas in Greece so that it effectively supports the recovery of the species in the country.

## Data Accessibility Statement

All data and codes used to carry out the analyses and generate the maps are provided as Supplementary Material (Appendix 3).

Additional Supporting Information may be found in the online version of this article:

**Appendix S1**: Details on the methods

**Appendix S2**: Predictors data (Bear_backg_env2017.csv); Observations data (Bear_spxy2017.csv); Habitat data (Habitat table, N2k raw, Tabke N2k, Table N2k_final)

